# Novel, active, and uncultured hydrocarbon degrading microbes in the ocean

**DOI:** 10.1101/2024.01.20.576437

**Authors:** Kathryn L. Howe, Julian Zaugg, Olivia U. Mason

**Author notes:** Address correspondence to Olivia U. Mason.

## Abstract

Given the vast quantity of oil and gas input to the marine environment annually, hydrocarbon degradation by marine microorganisms is an essential ecosystem service. Linkages between taxonomy and hydrocarbon degradation capabilities are largely based on cultivation studies, leaving a knowledge gap regarding the intrinsic ability of uncultured marine microbes to degrade hydrocarbons. To address this knowledge gap, metagenomic sequence data from the Deepwater Horizon (DWH) oil spill deep-sea plume was assembled to which metagenomic and metatranscriptomic reads were mapped. Assembly and binning produced new DWH metagenome assembled genomes that were evaluated along with their close relatives, all of which are from the marine environment (38 total). These analyses revealed globally distributed hydrocarbon degrading microbes with clade specific substrate degradation potentials that have not been reported previously. For example, methane oxidation capabilities were identified in all *Cycloclasticus*. Further, all *Bermanella* encoded and expressed genes for non-gaseous n-alkane degradation; however, DWH *Bermanella* encoded alkane hydroxylase, not alkane 1-monooxygenase. All but one previously unrecognized DWH plume members in the SAR324 and UBA11654 coded for aromatic hydrocarbon degradation. In contrast, *Colwellia* were diverse in the hydrocarbon substrates they could degrade. All clades encoded nutrient acquisition strategies and response to cold temperatures, while sensory and acquisition capabilities were clade specific. These novel insights regarding hydrocarbon degradation by uncultured planktonic microbes provided missing data, allowing for better prediction of the fate of oil and gas when hydrocarbons are input to the ocean, leading to a greater understanding of the ecological consequences to the marine environment.

**Importance:** Microbial degradation of hydrocarbons is a critically important process promoting ecosystem health, yet much of what is known about this process is based on physiological experiments with a few hydrocarbon substrates and cultured microbes. Thus, the ability to degrade the diversity of hydrocarbons that comprise oil and gas by microbes in the environment, particularly in the ocean, is not well characterized. Therefore, this study aimed to utilize non-cultivation based ‘omics’ data to explore novel genomes of uncultured marine microbes involved in degradation of oil and gas. Analyses of newly assembled metagenomic data and metagenomic and metatranscriptomic read recruitment revealed globally distributed hydrocarbon degrading marine microbes with clade specific substrate degradation potentials that have not been previously reported. This new understanding of oil and gas degradation by uncultured marine microbes suggested that the global ocean harbors a diversity of hydrocarbon degrading Bacteria, that can act as primary agents regulating ecosystem health.

## Introduction

Annually, hundreds of millions of liters of oil is introduced to the ocean (Head, et al. 2006), raising global concerns regarding the adverse effects of oil pollution on biology and overall marine ecosystem function (Brussaard et al. 2016). This oil originates from both natural and anthropogenic sources and is sufficient to cover the world ocean in a thin layer of oil (Head et al. 2006). The ocean is also a natural source of gaseous hydrocarbons, such as methane, a potent greenhouse gas. It is estimated that the ocean contributes 1–3% of global atmospheric methane (Saunois et al., 2020). Thus, the consumption of oil and gas by microbes in the water column is an important ecosystem service, both in terms of marine environmental health and for the capture of greenhouse gases that would otherwise efflux to the atmosphere.

One such example of both the magnitude of oil and gas that can be input to the marine environment, and the important role that microbes play in consuming hydrocarbons, is the Deepwater Horizon (DWH) oil spill. From April to July 2010 nearly 5.3 x 10^11^ g of oil and 1.7 x 10^11^ g of natural gas were input to the Gulf of Mexico (GOM) water column (Reddy et al., 2011). The depth at which the spill occurred (1,500 mbsl) prevented any photo-oxidation or oil recovery from the deep-sea water column. Therefore, microbial biodegradation of hydrocarbons was the primary mitigation taking place during the DWH spill in plume (Kujawinski et al. 2020). Thus, the role of indigenous deep-sea microbes in oil and gas consumption is paramount in terms of bioremediation and the environmental impact of deep-sea oil spills, such as the DWH spill in 2010, and marine oil spills in general.

Although there are few studies of the planktonic microbes in the deep-sea water column, analysis of uncontaminated deep-sea samples revealed a diversity of microbes (Mason et al., 2012; Redmond & Valentine, 2012), the majority of which are uncultured and, therefore, have unknown physiologies. This diversity decreased in seawater contaminated with DWH hydrocarbons, relative to uncontaminated samples (Mason et al. 2012; Rivers et al. 2013). The shifts in the taxonomic composition and abundance of indigenous deep-sea microbiota that led to decreasing diversity is well documented (Hazen et al. 2010; Valentine et al. 2010; Kessler et al. 2011; Redmond and Valentine 2012; Mason et al. 2012; Dubinsky et al. 2013; Kleindienst et al. 2016). These studies revealed a pattern in succession with *Oceanospirillales* (now recognized to be *Bermanella* and will be referred to as such for the metagenome assembled genomes (MAG) presented herein) dominating early in the spill followed by *Colwellia*, *Cycloclasticus* and finally by methylotrophs. The changes in the dominant microbial players was attributed to variation in the supply of different hydrocarbons during the spill (Dubinsky et al., 2013). For example, *Oceanospirillales* dominated during the period of unrestricted flow, when the concentration of n-alkanes and cycloalkanes was highest (Dubinsky et al., 2013). On June 5^th^, partial capture of hydrocarbons led to benzene, toluene, ethylbenzene and xylene (BTEX) and natural gases dominating the plume (Reddy et al., 2011), which coincided with an increase in abundance of *Colwellia* and *Cycloclasticus* (Dubinsky et al., 2013).

Although more than a decade has passed since the DWH spill, an information gap regarding microbial hydrocarbon degradation persists, which diminishes our ability to understand the ecosystem effects of both past and future oil spills in the marine environment. For example, although gaseous hydrocarbons were the most abundant hydrocarbon input during the spill (Reddy et al., 2011), and evidence that oxidation of methane (and ethane and propane (Valentine et al., 2010)) was an active microbial process (Kessler et al. 2011; Crespo-Medina et al. 2014), methane oxidation capabilities remain largely unattributed to any specific microbe (Kujawinski et al. 2020). A variety of approaches including particulate methane monooxygenase (*pmo*) gene phylogenetic reconstruction (Crespo-Medina et al., 2014), stable isotope probing (SIP) coupled with 16S rRNA gene sequencing (Redmond & Valentine, 2012), metagenomic (Mason et al., 2012) and metatranscriptomic sequence annotation (Mason et al. 2012; Rivers et al. 2013) revealed a diversity of *pmo* genes and transcripts, but taxonomy and function of indigenous methane oxidizing bacteria was not definitively linked through genome assembly.

Non-gaseous straight chain n-alkanes, such as n-pentane and n-hexane, the cyclic alkanes cyclopentane and cyclohexane, and BTEX aromatics were also abundant (Ryerson et al., 2011, 2012). Polycyclic aromatic hydrocarbons (PAHs) were observed in the plume at levels that would have acute toxicity effects (Diercks et al., 2010), at the surface (Hazen et al. 2010), and on the seafloor (Mason et al. 2014). The degree of resolution on microbial consumption of non-gaseous hydrocarbons varies. For example, single amplified genome (SAG) recovery revealed n-alkane and cycloalkane degradation capabilities in the plume which was ascribed to indigenous *Oceanospirillales* (Mason et al., 2012). This *Oceanospirillales* SAG has genes coding for alkane monooxygenase (*alkB*), cyclohexanol dehydrogenase and cyclohexanone monooxygenase (Mason et al., 2012). However, the 16S rRNA gene encoded in the SAG was 95% similar to the dominant indigenous *Oceanospirillales* and the SAG recruited a low number of DNA and RNA reads (Mason et al., 2012). Therefore, how well it represented the hydrocarbon degradation pathways in the dominant *Oceanospirillales* in the plume is not clear.

Even less is known about what plume microbes were capable of degrading aromatics. Culturing experiments have revealed that *Cycloclasticus* can grow on aromatics (Head et al. 2006), including toluene (Dyksterhouse et al., 1995; Wang et al. 1996), xylene (Wang et al., 1996) and PAHs (Geiselbrecht et al., 1996, 1998). SIP experiments revealed a possible role for *Colwellia* in degrading benzene (Redmond & Valentine, 2012); however, no further resolution has been provided for what indigenous microbes could degrade aromatics in the deep-sea.

Thus, much remains unknown regarding the degradation capabilities of uncultured, indigenous deep-sea microbes more than a decade since the DWH oil spill, during which time new and improved metagenomic assembly and classification tools have become available. We have taken advantage of such tools to assemble the metagenomic data presented in Mason et al. (2012), obtaining 12 DWH MAGs of medium to high quality (Bowers et al., 2017). Previous attempts to assemble this data were largely unsuccessful (Mason et al. 2012), thus important information regarding degradation potential and identity remain unresolved. To expand our understanding of hydrocarbon degradation, nutrient acquisition, chemotaxis and motility, beyond the GOM to the global ocean, these DWH MAGs were analyzed together with 26 close relatives. This analysis provided missing linkages between taxonomy coupled with function, which previously hampered our understanding of microbial hydrocarbon oxidation in the marine environment, thus minimizing our ability to predict the ecological consequences of hydrocarbons input to the ocean.

## Materials and Methods

### Sample Collection

As described in Mason et al. (2012) 15 L of seawater were filtered through a 0.2 µm diameter filters from stations sampled in the Gulf of Mexico during two cruises from May 27-June 2 2010 on the *R/V Ocean Veritas* and *R/V Brooks McCall*. Details regarding sample collection were provided in Hazen et al. (2010).

### DNA and RNA extraction, processing, and sequencing

A detailed extraction protocol was provided in Mason et al. (2012). Briefly, DNA was extracted from microbial cells collected onto filters using a modified Miller method (Miller et al. 1999), with the addition of a pressure lysis step to increase cell lysis efficiency. Sample material was transferred to a Lysing Matrix E tube (MP Biomedicals, Solon, OH) and the samples were subjected to bead beating, followed by centrifugation at which time chloroform was added to the supernatant. After centrifugation the aqueous phase clean-up procedures followed the instructions in the MoBio Soil DNA extraction kit.

Total RNA was processed as described in Mason et al. (2012). In summary, total RNA from the proximal and distal plume stations was amplified, followed by first strand synthesis of cDNA and double stranded cDNA. Poly(A) tails were removed from purified cDNA by digesting purified DNA with *Bpm*I for 3 h at 37°C. Digested cDNA was subsequently purified.

To increase yields required for sequencing, DNA and cDNA were amplified by emulsion PCR. A detailed description of this method can be found in (Blow et al., 2008). Amplified DNA was sequenced using the Illumina GAIIx with 2 x 114 bp pair-end technology, resulting in 14–17 Gb of sequence read data per sample. These reads are publicly available in NCBI (PRJNA336903-05) in IMG Gold Study ID Gs0063184 and on the Mason server http://mason.eoas.fsu.edu in the DWH_plume directory. Amplified cDNA was sequenced using the Illumina GAIIx sequencing platform with 2x100 bp pair-end technology. These reads are publicly available in NCBI (PRJNA839076). MAGs are also available in NCBI under SAMN37094393-95, SAMN37094224-28, SAMN37094233-36 and are linked with their respective metagenome sequence data in NCBI (PRJNA336903-05).

### Metagenomic sequencing, assembly, annotation and read mapping

The quality of paired-end metagenomic reads was assessed, and adapter sequences identified, using FastQC (ver. 0.10.1, http://www.bioinformatics.babraham.ac.uk/projects/fastqc/). Low quality reads and adapters were subsequently removed using Trimmomatic (ver. 0.36, Bolger, et al., 2014, ILLUMINACLIP:TrueSeq and over represented adapters:2:30:10, LEADING:3, TRAILING:3, SLIDINGWINDOW:4:15, MINLEN:50). Those reads lacking a mate pair were removed. Reads were assembled using MetaSPAdes (ver. 3.13, Nurk, et al., 2017) with default parameters. Contigs less than 500bp were removed using BBMap (ver. 38.35, https://sourceforge.net/projects/bbmap/). Filtered reads were mapped onto their respective scaffolds using CoverM ‘make’ (ver. 0.2.0, https://github.com/wwood/CoverM), with CoverM ‘filter’ used to remove low quality mappings (minimum identity 95% and minimum aligned length of 50bp). Scaffolds for each sample were binned using UniteM (ver. 0.0.15, https://github.com/dparks1134/UniteM), with a minimum contig length of 1,500bp and Maxbin (ver. 2.2.4) (Wu et al., 2014), MetaBAT (ver. 0.32.5) (Kang et al., 2015) and MetaBAT2 (ver. 2.12.1) (Kang et al., 2019) binning methods (max40, max107, mb2, mb_verysensitive, mb_sensitive, mb_specific, mb_veryspecific and mb_superspecific). From this assembly approach 12 DWH MAGs were obtained (Table 1). The completeness and contamination of all bins were calculated by CheckM (ver. 1.0.12) (Parks et al., 2015). These MAGs were medium (> 50% complete and < 10% contamination) and high quality (> 90% complete and < 5% contamination), except the three *Colwellia*, which met one of the two criteria (Table 1). Taxonomy was assigned using the Genome Taxonomy Toolkit (GTDB-Tk, ver. 2.3.0; with reference to GTDB R08-RS214) (Chaumeil et al., 2019) with up to 120 bacterial single-copy marker proteins, which are predominantly ribosomal (see Supp. Table 6 from Parks et al. 2017). The GTDB-Tk phylogenetic tree based on the alignment of concatenated single-copy marker proteins was viewed using Archaeopteryx (Han & Zmasek, 2009) to identify close relatives of the MAGs using phylogeny with monophyly and branch length as decision making criteria for inclusion (see Supp. Figure 1). These 26 additional genomes were included in all subsequent analyses.

**Table 1.**
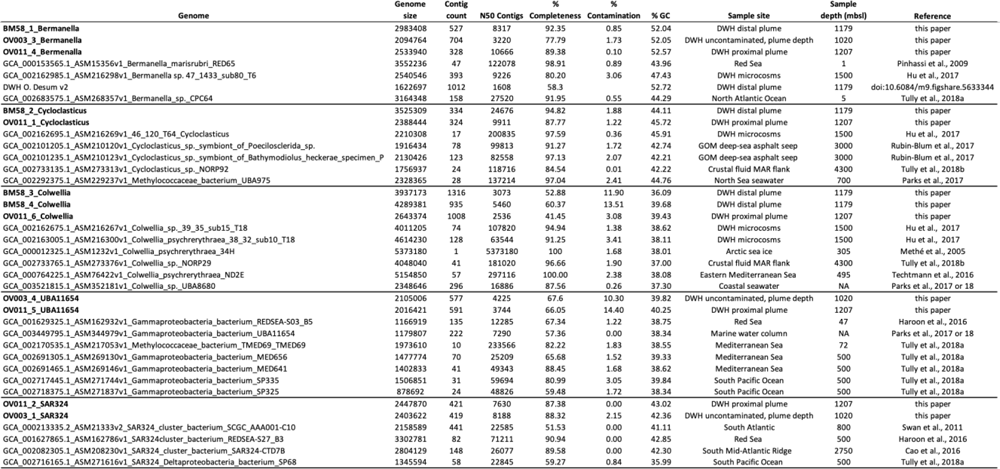
Detailed information regarding genome statistics, including completeness and contamination percentages, sampling locations and depths.

All genomes were analyzed using Anvi’o (ver. 6.2) (Eren et al. 2015). In Anvi’o, Prodigal was used to identify open reading frames (ORFs). ORFs were annotated with ‘anvi-run-pfams’ and ‘anvi-run-ncbi-cogs’, based on the Pfam (Bateman et al., 2004) and NCBI’s Clusters of Orthologous Genes (COG) (Tatusov et al., 2000) databases, respectively. Pfam and COG annotations were also verified using blastx with DIAMOND (ver. 0.9.30) (Buchfink et al. 2014) to compare ORFs to NCBI’s non-redundant RefSeq protein dataset (accessed on 03/10/2020) (Tatusova et al., 2016). Functional annotations were confirmed when two of three databases (COGs, Pfams, and RefSeq) agreed.

To determine genome, gene and transcript abundances MG and MT reads were mapped to DWH MAGs and non-DWH genomes using Bowtie 2 (ver. 2.3.4.1) (Langmead & Salzberg, 2012) following the script provided at https://merenlab.org/data/tara-oceans-mags/ and used in Thrash et al. (2017) and more recently as the RRAP pipeline (Kojima et al., 2022). Before mapping cDNA reads ribosomal RNA was subtracted using riboPicker (ver 0.4.3) with the default settings (Schmieder et al. 2012). Low quality read mappings were filtered using BamM (ver. 1.7.3, http://ecogenomics.github.io/BamM/) with --percentage_id 0.95 --percentage_aln 0.75. Alignment summary statistics were obtained using SAMtools (ver. 1.7-2) (Li et al., 2009) command idxstats with default settings. Subsequently, genome, gene and transcript abundances were normalized by calculating reads per kilobase per million mapped (RPKM) values according to Mortazavi et al. (2008), and detailed in Thrash et al. (2017) and Kojima et al. (2022). RPKM values are reported as MG RPKM maximum/MT RPKM maximum in the results. Following normalization, average RPKM values were calculated, with low values being below average and high values being above average.

### Pangenomic analysis

A comparative genomic analysis of the 12 DWH MAGs and 26 non-DWH genomes was performed using the Anvi’o pangenomic workflow v2 (Eren et al., 2015). Hits between gene clusters, the representative sequences of predicted ORFs that are grouped together based on homology at the translated DNA sequence level, and bacterial single-copy core genes were identified using HMMER and hidden Markov models (anvi-run-hmms). The pangenome was generated using DIAMOND (Buchfink et al., 2014) as part of the ‘anvi-pan-genome’ program, which was run in sensitive mode with hierarchical clustering enforced. Core genes encoded in all 38 genomes were identified, as was a DWH pangenome for the core genes among only the 12 DWH MAGs. Average nucleotide identity (ANI) among all genomes was calculated using Sourmash with --containment -- ani --ksize 31 (Brown and Irber 2016).

## Results

### DWH MAG assembly, taxonomy and close relatives from the global ocean

Twelve MAGs were assembled from the DWH proximal plume (OV011), distal plume (BM58) and from a non-oil contaminated sample collected at plume depth (OV003) from metagenomes that were presented in Mason et al (2012) but were not assembled. These MAGs represent four different clades in the Gammaproteobacteria and SAR324 (formerly part of the Deltaproteobacteria, now a candidate phylum (Parks et al., 2018)). The new DWH MAGs presented in the current study are classified as *Bermanella* (3 MAGs), *Colwellia* (3 MAGs), *Cycloclasticus*, (2 MAGs), UBA11654 and SAR324 (2 MAGs each) (Table 1). These MAGs averaged 75% completion and had low contamination (avg. 5%) (Table 1).

The DWH MAGs were evaluated alongside non-DWH genomes that were medium to high quality (Table 1, average of 83% completion and 1.36% contamination. While the DWH MAGs were most similar to one another, phylogenetic analysis revealed non-DWH bacteria sampled from the global ocean (Table 1) were also closely related. For example, as shown in Table 1, close relatives were sampled in the Red Sea (Haroon et al., 2016; Pinhassi et al., 2009), the North Atlantic Ocean (Tully et al. 2018a), South Pacific Ocean (Tully et al. 2018a), Mediterranean Sea (Techtmann et al., 2016; Tully et al., 2017), the North Sea (Parks et al., 2017), and also from crustal fluids collected on the flanks of the Mid-Atlantic Ridge (Tully et al. 2018b), symbionts of mussels and sponges living at hydrocarbon seeps (Rubin-Blum et al., 2017), and microcosms with Macondo oil (Hu et al. 2017).

Average nucleotide identity (ANI) calculations showed DWH MAGs had the highest ANI with other DWH MAGs within the same clades (Fig 1). However, there were several other genomes that also had high ANI with DWH MAGs. Most notably, the “DWH O. Desum v2” MAG (Eren, 2017), which was previously assembled from the metagenome data presented in Mason et al. (2012), had > 99% ANI to the *Bermanella* DWH MAGs (Fig 1). MAGs described in Hu et al. (2017) that were obtained from microcosms with Macondo oil were 70% to more than 90% similar to DWH *Bermanella*, *Cycloclasticus* and *Colwellia* MAGs (Fig 1), although DWH *Colwellia* MAGs were equally similar to *Colwellia psychrerythraea 34H*, isolated from Artic sea ice (Methé et al., 2005). UBA11654 DWH MAGs were most similar to each other at 98%, with 75% or less in similarity with non-DWH UBA11654 genomes (Fig 1). The SAR324 DWH MAGs were most similar to each other and two non-DWH SAR324 MAGs (Fig 1).

**Figure 1.**
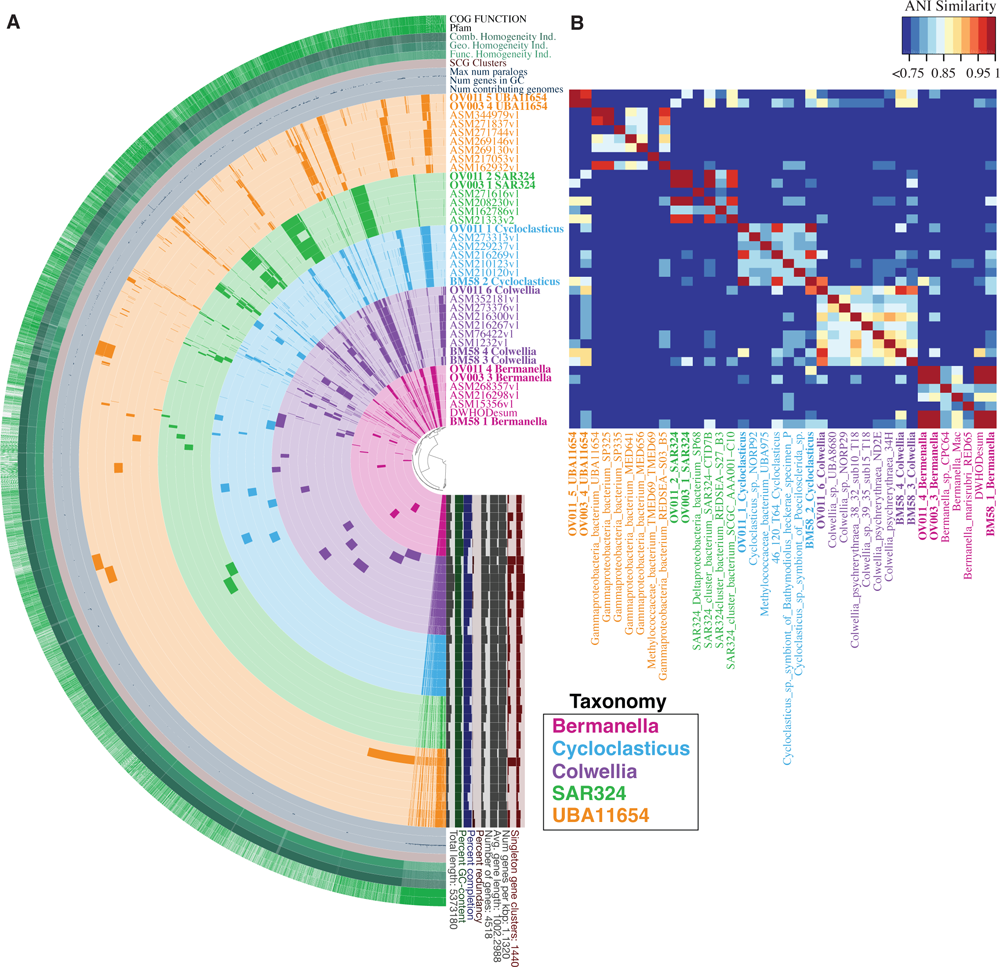
Phylogram (A) and ANI plot (B) of all 38 MAGs and genomes analyzed. ANI plot (2B) only shows ANI above 75%. Genome names are color coded by taxonomy with DWH MAGs in bold text.

### Microbial abundances and transcriptional activity

Microbial abundances and transcriptional activity were determined by mapping unassembled MG and MT reads from Mason et al. (2012) to the 38 DWH and non-DWH genomes included in this study. Read counts were normalized by determining reads per kilobase per million (RPKM) and are reported as low if RPKM values were below average, or high if RPKM values were above average. DWH *Bermanella* MAGs had the highest RPKM abundances, ranging from 67 in the distal plume to 55 in the proximal plume MG (Fig 2), compared to an average RPKM abundance of 5. These MAGs also recruited the highest number of MT reads of any genomes, with the highest abundance (RPKM of 32) observed in the proximal plume and up a to an RPKM of 12 in the distal plume (Fig 2), which was high compared to the averaged genome abundance of RPKM 1. A large number of MG reads also mapped to DWH O. Desum v2 (RPKM ranged from 47–53), but was not highly represented when recruiting MT reads (RPKM abundances were 0.26 and 6.6) (Fig 2). While less abundant, MG and MT reads did map to all other *Bermanella* genomes (Fig 2).

**Figure 2.**
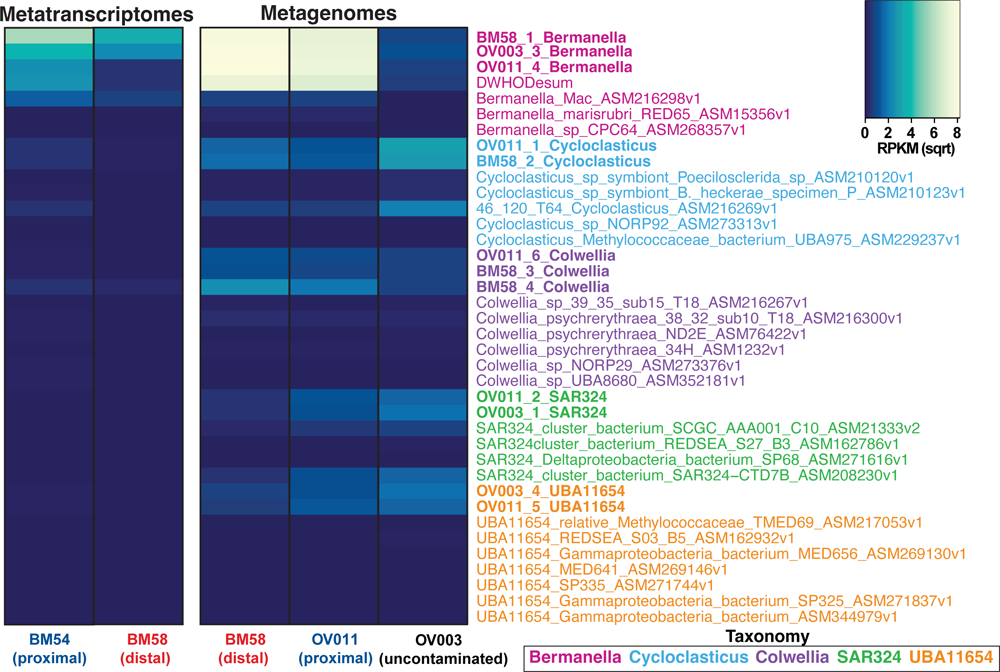
Microbial abundance and activity determined read recruitment of the MGs and MTs to the 38 MAGs and genomes in this study. Genome names are color coded by taxonomy with DWH MAGs in bold text.

DWH *Cycloclasticus* and *Colwellia* MAGs recruited MG reads and low levels of MT reads (Fig 2). Specifically, DWH *Cycloclasticus* MG RPKM abundances ranged from 2–9, with the greatest number of reads recruited in the uncontaminated sample collected from plume depth (Fig 2), while DWH *Colwellia* had MG RPKM abundances ranging from 0.73 up to 6 (Fig 2). These MAGs recruited MT reads, but RPKM values were low in comparison to *Bermanella* (Fig 2). Non-DWH *Cycloclasticus* and *Colwellia* genomes recruited a below average amount of MG and MT reads relative to other genomes (Fig 2). Finally, UBA11654 and SAR324 DWH MAGs read recruitment ranged from RPKM of 0.20–3.22 (MG) to < 0.005 (MT) (Fig 2). All non-DWH UBA11654 and SAR324 MAGs recruited MG reads. All but one SAR324 genome recruited MT reads from a plume sample, albeit at low levels relative to other genomes, while no UBA11654 genomes recruited any MT reads (Fig 2).

### Hydrocarbon degradation

### Gaseous hydrocarbons

All *Cycloclasticus* genomes, including DWH *Cycloclasticus* MAGs, coded for methane oxidation (Figs 3 & 4). Outside of the *Cycloclasticus,* only a DWH *Colwellia* MAG encoded methane oxidation (Figs 3 & 4). Specifically, all *Cycloclasticus* and one *Colwellia* encoded methane/ammonia monooxygenase, subunits A–C (Fig 4). Only these DWH MAGs and one Macondo oil microcosm MAG *pmoABC* genes recruited MG and MT reads (Fig 4). From the uncontaminated deep-sea sample, DWH *Cycloclasticus* and *Colwellia pmoABC* genes recruited the most reads of any genes involved in hydrocarbon degradation (MG RPKM maximum was 15; Fig 4), which was well above the average of RPKM 5. In the plume samples, DWH *Cycloclasticus pmoABC* had the highest MG RPKM values in proximal and distal plumes followed by DWH *Cycloclasticus* and the DWH *Colwellia* (maximum RPKM was 2.3; Fig 4). DWH *Colwellia* from the proximal plume and DWH *Cycloclasticus* from the distal plume recruited the greatest number of MT reads to *pmoABC*, with a maximum RPKM of 1.38, which was the same as the average (Fig 4).

**Figure 3.**
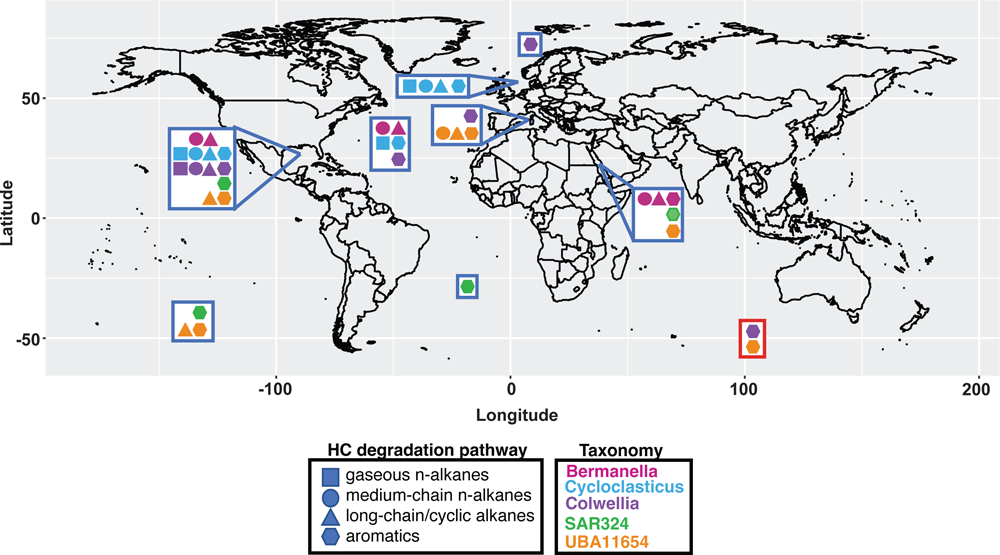
Global distribution of sample locations of genomes analyzed, with their hydrocarbon degradation capabilities shown by shape and color-coded by taxonomy. The red box encloses genomes that were assembled from the marine environment, but the exact location was not provided in Parks et al. (2017).

**Figure 4.**
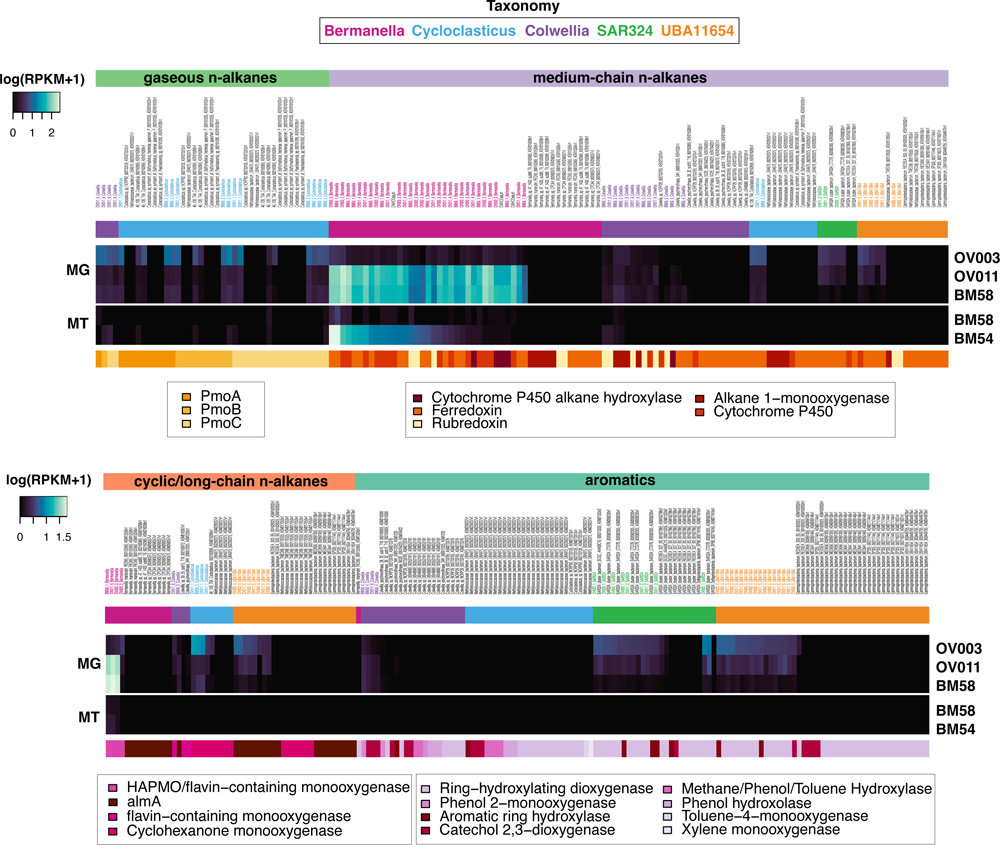
Heatmap of abundance (metagenome, MG) and expression (metatranscriptome, MT) of genes involved in hydrocarbon degradation. DWH MAG names are in bold text and color coded by taxonomy. Non-DWH genomes are in black text.

All genomes that recruited reads to *pmoABC* genes also encoded additional steps culminating in a partial methane oxidation pathway, including a short-chain alcohol dehydrogenase (SDR) (MG RPKM maximum of 193/MT RPKM maximum of 1.2), generic aldehyde dehydrogenase (ALDH) (37/0.33). Additionally, multiple formaldehyde oxidation pathways were encoded, with the tetrahydrofolate (H4F) pathway having the highest MG recruitment (RPKM maximum was 54; Supp. Fig 2.), while no MT reads were recruited (Supp. Fig 2). Of the remaining genomes that encoded *pmoABC,* but lacked read recruitment (four *Cycloclasticus*), two encoded a partial pathway while the other two encoded a full pathway including formate dehydrogenase (FDH). However, the only genes that recruited any MG reads were for the GSH pathway belonging to *Cycloclasticus sp. Symbiont of Bathymodiolus heckerae specimen P* (MG RPKM maximum of 0.17) and none of these genes recruited MT reads (Supp. Fig 2).

### Non-gaseous n-alkanes

All *Bermanella* encoded medium-chain (C_5_-C_26_) n-alkane degradation, but there was more than one pathway observed. For example, all DWH *Bermanella* MAGs and DWH O. Desum v2, which was assembled from our metagenomes and presented in Eren, 2017, coded for genes in the cytochrome P450 family, not alkane-1 monooxygenase (AlkB), an enzyme used by *Alcanivorax borkumensis* in the degradation of medium chain n-alkanes. Specifically, a cytochrome P450 alkane hydroxylase (CYP153) was encoded in all DWH *Bermanella* MAGs and DWH O. Desum v2. These genes were highly abundant in the plume, with MG RPKM values up to 54 (Fig 4). While present in the uncontaminated sample, RPKM values were low (below average) relative to the plume samples (Fig 4). All DWH *Bermanella* MAGs CYP153 genes were expressed in the plume samples, as was DWH O. Desum v2 (RPKM values for all were < 0.50; Fig 4). The CYP153 genes in the *Colwellia* DWH MAG BM58_3 had MG RPKM values of < 0.52 in the distal and proximal plumes, but was not detected in the uncontaminated sample, nor was it expressed (Fig 4). Three DWH *Bermanella* MAGs and DWH O. Desum v2 that encoded CYP153 also encoded a generic cytochrome P450 (not annotated as alkane hydroxylase) and ferredoxin. These genes recruited up to 184/263 (Fig 4). Three of these *Bermanella* genomes also encoded ferredoxin reductase, suggesting a three domain CYP153 architecture (Nie et al., 2014) for non-gaseous n-alkane degradation. These ferredoxin reductases genes recruited reads up to 59/< 2.

In contrast to DWH *Bermanella*, *alkB* was encoded in the genomes of all non-DWH *Bermanella*, in two DWH *Colwellia* MAGs, one non-DWH *Cycloclasticus* and one UBA11654 (Fig 4). DWH MAG *alkB* genes recruited MG reads from the proximal, distal plume and the uncontaminated sample (RPKM maximum was 1.82), while *Bermanella sp.* 47_1433_sub80_T6 (Hu et al. 2017) recruited a low level of reads (MG RPKM was 0.05) and only from the distal plume. The only actively expressed *alkB* genes were from a *Cowellia* DWH MAG, which recruited MT reads from both plume samples (RPKM maximum was 0.38; Fig 4).

AlkB is part of the two domain architecture (Nie et al., 2014) with rubredoxin mediating the second step in alkane oxidation (Sabirova et al. 2006). Although no DWH *Bermanella* MAGs encoded *alkB*, all had rubredoxin genes, as did *Bermanella marisrubri RED65* and *Bermanella sp. CPC64*, but not DWH O. Desum v2 or *Bermanella sp. 47 1433 sub80 T6*. DWH *Bermanella* rubredoxin genes were abundant in the proximal and distal plume (RPKM maximum was 14), but less so in the uncontaminated sample (RPKM maximum was 0.40; Fig 4). These genes were expressed in both plume samples with RPKM values up to 14 (Fig 4). The remaining *Bermanella* MAGs encoding rubredoxin genes did not recruit any MG or MT reads (Fig 4). The two DWH *Colwellia* MAGs that had *alkB* genes also encoded rubredoxin, but recruited relatively few MG and MT reads (all RPKM < 1) compared to DWH *Bermanella* (Fig 4).

### Long-chain n-alkanes and cyclic alkanes

There was a diversity of genes encoding long-chain n-alkane (>C_18_) and cyclic alkane degradation in all clades analyzed here, except SAR324. In particular, this was a ubiquitous pathway in DWH and non-DWH *Bermanella*. For example, all DWH *Bermanella*, including DWH O. Desum v2, and two non-DWH *Bermanella* coded for genes that were most similar (58%) to a 4-hydroxyacetophenone monooxygenase (HAPMO) gene in a MAG from a subseafloor aquifer (Tully et al 2018b). These genes were equally as similar to non-HAPMO enzymes annotated as flavin-containing monooxygenases. We also would note that these genes have low sequence similarity with *almA* and with *A. borkumensis’* cyclohexanone monooxygenase (30% and 24%, respectively). Overall, these flavin containing monooxygenases were highly abundant with MG RPKM abundances up to 58 in plume samples, but were less abundant in the uncontaminated sample (MG RPKM maximum was 10) and expressed at low levels in the plume samples (MT RPKM maximum was 0.72; Fig 4). Of the long-chain and cyclic alkane gene annotations, those encoded by the aforementioned *Bermanella* were the only genes that were expressed (Fig 4).

Other flavin-containing monooxygenases were identified in DWH and non-DWH *Bermanella, Cycloclasticus,* and UBA11654. These flavin-binding monooxygenase genes were most abundant in the uncontaminated sample (MG RPKM maximum was 9), compared to the plume samples (MG RPKM maximum was 1.7; Fig 4). None of these genes were expressed (Fig 4). In this group, DWH and non-DWH *Cycloclasticus* and UBA11654 genes and non-DWH *Bermanella* were annotated as *almA.* The *almA* enzyme is a well-known flavin-binding monooxygenase, that functions as an alkane hydroxylase in *A. hongdengensis* and *dieselolei* allowing for microbial degradation of long-chain n-alkanes (C. Liu et al., 2011; W. Wang & Shao, 2012).

Finally, cyclohexanone monooxygenases, a BVMO involved in degradation of cyclic alkanes, was identified in DWH and non-DWH *Cycloclasticus* and non-DWH UBA11654. Only DWH *Cycloclasticus* cyclohexanone monooxygenases recruited reads, with a MG maximum RPKM of 10 in the uncontaminated sample and RPKM values of < 1 in the plume sample (Fig 4). None of these genes recruited MT reads (Fig 4).

### Aromatics

Members of all clades, including one non-DWH *Bermanella*, encoded for aromatic hydrocarbon degradation genes which were primarily ring-hydroxylating dioxygenases, with a lower number of catechol 2,3-dioxygenases. DWH and non-DWH members of *Colwellia*, UBA11654, and SAR324 were the only clades to recruit any reads to these genes, with the largest number of MG reads from the distal plume (station BM58), although RPKM values were low compared to other hydrocarbon degradation pathways (maximum RPKM was 1.3; Fig 4). A non-DWH SAR324 MAG (SAR324 cluster bacterium SAR324-CTD7B) recruited the greatest number of MG reads, with a maximum RPKM of 3.3 from the proximal plume (Fig 4). Despite the abundance of BTEX hydrocarbons in the plume (Reddy et al., 2011), only two genes specifically coding for BTEX degradation, toluene-4 monooxygenase and xylene monooxygenase, were observed in any genome, and only in the non-DWH *Cycloclasticus Methylococcaceae bacterium UBA975* (Parks et al., 2017), which did not recruit any MG or MT reads (Fig 4). Overall, RPKM values were highest when recruiting MG reads from the uncontaminated, plume depth station (OV003) to genes coding for aromatic hydrocarbon degradation identified in MAGs, with non-DWH and DWH SAR324 and UBA11654 genes recruiting the greatest number of reads (maximum MG RPKM was 6.4; Fig 4). None of the aromatic hydrocarbon degrading genes encoded in the genomes analyzed here recruited any transcripts (Fig 4).

#### Nitrogen, sulfur, phosphorus and iron

In evaluating the capacity for carrying out anaerobic respiration through denitrification, most *Colwellia* MAGs and genomes in this study encoded the necessary enzymes to carry out complete denitrification (Fig 5). Specifically, DWH and non-DWH *Colwellia* nitrate and nitrite reductase genes recruited a maximum of 58/3, both well above the averages of 6.8/0.3, with lower recruitment to nitric oxide reductase (all RPKM < 3) and nitrous oxide reductase (all RPKM < 1; Fig 5). Most UBA11654 MAGs encoded both nitrate and nitric oxide reductases (MG and MT RPKM < 4), but not nitrite or nitrous oxide reductases (Fig 5). DWH and non-DWH *Bermanella* and *Cycloclasticus* MAGs encoded nitrite reductase with RPKM values of 40–60, but no other steps in the denitrification pathway. Most SAR324 encoded nitric oxide reductase with MG read recruitment RPKM < 3 across all samples. Beyond denitrification, all clades encoded for nitrate/nitrite transporters, with one DWH *Colwellia* and DWH O. Desum v2 recruiting the greatest number of MG reads (RPKM > 50; Fig 5). DWH *Bermanella* and a DWH UBA11654 nitrate/nitrite transporters recruited the highest number of MT reads (RPKM 12–15; Fig 5).

**Figure 5.**
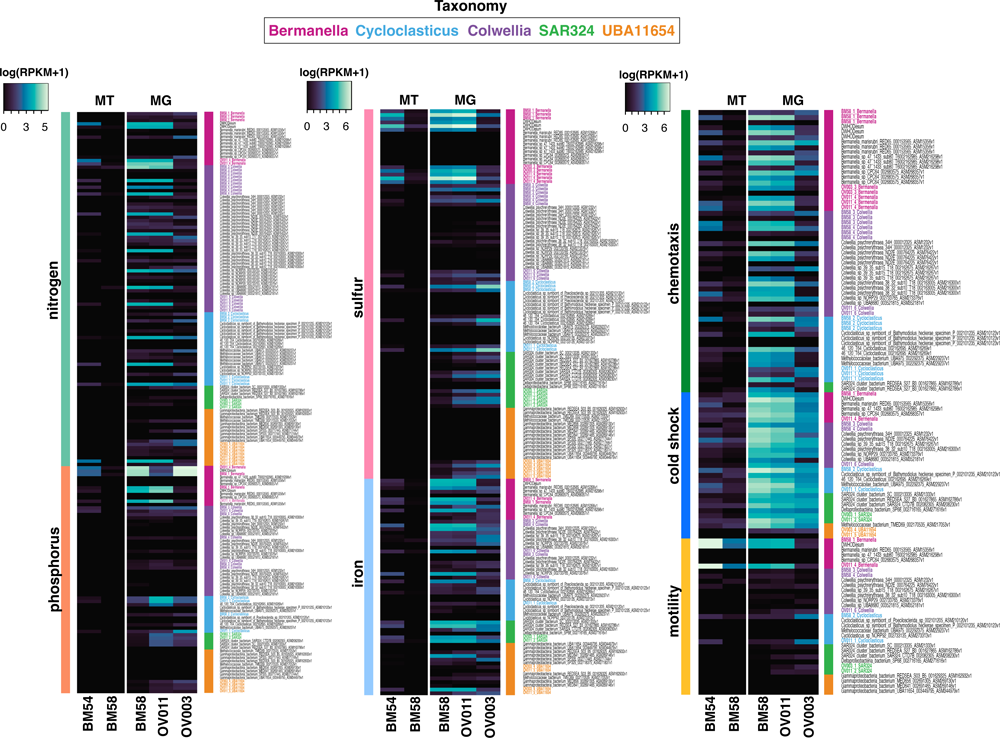
Heatmap of abundance and expression of genes involved in anaerobic respiration, nutrient acquisition, chemotaxis, motility, and cold-shock. DWH MAG names are in bold text and color coded by taxonomy. Non-DWH genomes are in black text.

Genome annotation did not reveal pathways for dissimilatory sulfate reduction, nor sulfur oxidation. However, genes coding for assimilatory sulfate reduction, including ATP sulfurylase, adenylyl sulfate kinase, 3’-phosphoadenosine 5’-phosphosulfate reductase, and sulfite reductase, encoded by DWH *Bermanella* and O. Desum v2 recruited MG reads up to RPKM values of 69 with an average of 22 and MT reads up to RPKM values of 42 with an average of 2.5 (Fig 5). All non-DWH genomes and three DWH MAGs coded for assimilatory sulfate reduction, but these genes recruited low levels of MG and MT reads (RPKM maximum of 0.2). Sulfur transport genes encoded by DWH *Bermanella* recruited MG reads up to an RPKM maximum of 981 and MT up to a PRKM maximum of 163 from the plume samples (Fig 5). Both DWH *Cyclcolasticus* MAGs and on non-DWH *Cyclcolasticus* recruited MG reads from the uncontaminated sample (RPKM maximum was 286), but not from plume samples. The remaining genomes had MG and MT RPKM values of < 2 to sulfur transport genes (Fig 5).

Phosphate cycling genes include alkaline phosphatase (APase), utilized in organophosphate acquisition, and phosphoenolpyruvate (PEP) phosphomutase, which is involved in phosphonate metabolism, with RPKM averages of 11/0.28 were identified. APase was encoded by members of most clades, with DWH *Bermanella* MAGs recruiting the most MG reads (RPKM max was 120) of the microbes analyzed and was the sole clade that recruited MT reads (RPKM < 1) to these genes (Fig 5). PEP phosphomutase was also encoded by all clades, except SAR324, with MG and MT read recruitment of RPKM < 3 (Fig 5). No other phosphonate metabolism genes were annotated in any genome (Fig 5). All clades had members that encoded phosphate transport genes (ABC type phosphate/phosphonate transport system) with DWH *Bermanella* recruiting the greatest number of MG reads (RPKM maximum was 363; Fig 5). These DWH *Bermanella* MAGs were the only microbes that recruited MT reads to phosphate transport genes with a maximum RPKM of 13 (Fig 5).

Siderophore production and transport genes were encoded by 29 genomes in all clades except SAR324. A DWH *Colwellia* siderophore-associated genes recruited the most MG (RPKM maximum was 132) and MT reads (RPKM maximum was 12) of any microbe in this study (Fig 5). Additionally, iron acquisition and transport genes (including ferrous and ferric iron transport systems) were encoded by members of all clades with RPKM averages of 18/0.6. These genes generally recruited a similar magnitude of MG reads in the proximal and distal samples for each genome, while recruiting lower MG reads in the uncontaminated sample (Fig 5). For example, the ferric iron transport gene of a non-DWH SAR324 genome had MG RPKM values of 143 and 148 in the plume samples, 23 in the uncontaminated sample, and MT RPKM maximum of 8 (Fig 5).

#### Chemotaxis and motility

Nearly all *Bermanella*, *Colwellia*, and *Cycloclasticus* encoded methylase, signaling, and receiver domains of methyl-accepting chemotaxis proteins (MCPs) and flagellar-associated proteins, indicating the capacity for chemotaxis (RPKM averages of 54/2) and motility (RPKM averages of 144/2.4). No UBA11654, and only one SAR324, encoded these genes. DWH *Bermanella* chemotaxis genes had the maximum MG RPKM of 394 and 321 from the distal and proximal station samples respectively, while motility genes from DWH *Cycloclasticus* recruited the most MG reads (684 and 413 in the distal and proximal sites, respectively; Fig 5). Fewer MG reads were recruited to chemotactic and motility genes in the uncontaminated sample compared to plume samples. These genes recruited up to MT RPKM of 17 for motility, 29 for chemotaxis (Fig 5).

#### Cold tolerance

Cold shock genes in the CspA family were encoded by members of all clades, with RPKM averages of 8/45. The cold shock genes of DWH *Bermanella* MAGs recruited high MG reads (RPKM > 150) in the distal and proximal samples (Fig 5). These same MAGs also recruited a large number of MT reads (RPKM > 890) in the proximal sample, but fewer reads in the distal (RPKM < 26; Fig 5). The remaining MAGs recruited fewer MG or MT reads (RPKM < 20; Fig 5).

#### The pangenome and clade core functions

Given the paucity of data available on the genomic content of deep-sea planktonic microorganisms, particularly those involved in hydrocarbon degradation, a core genome, or pangenome was identified for all 38 genomes and separately for DWH MAGs only. Analysis of all genomes (100, 262, 742 nucleotides) revealed that the core functions identified in the pangenome included acyl-CoA dehydrogenase, acetyl-CoA acetyltransferase, and enoyl-CoA hydratase/isomerase (Supp. Table 1). DWH *Bermanella* MAGs recruited the highest number of MG and MT reads to these genes with a maximum of 173/26. Short-chain alcohol dehydrogenase (SDR), utilized in methane oxidation and other metabolic pathways, would have been included with the pangenome, but the COG and Pfam annotations for these genes in one DWH *Bermanella* (OV003_3) were not consistent (as shown in Supp. Fig 2). The remaining core functions consisted mostly of housekeeping genes (Supp. Table 1). The core genes of just the DWH MAGs included the genes in the full pangenome core and additional genes associated with hydrocarbon degradation: citrate synthase and aldehyde dehydrogenase (Supp. Table 1).

## Discussion

The first hydrocarbon degrading microbes were cultured and described more than a century ago. In fact, much of what we know about hydrocarbon degrading microbes, particularly those in the marine environment, stems from cultivation-based studies of microbes such as *A. borkumensis*, an aliphatic hydrocarbon degrader that is ubiquitous in marine ecosystems and blooms quickly when hydrocarbons are input to the environment (Yakimov et al., 1998; Schneiker et al., 2006). Other Gammaproteobacteria such as *Cycloclasticus* (Dyksterhouse et al. 1995) and *Oleispira* (Yakimov et al., 2003) were also isolated from the marine environment, and, like *A. borkumensis* were shown to be obligate hydrocarbon degrading (hydrocarbonoclastic) bacteria. These early studies provided important linkages between hydrocarbon degradation capabilities and taxonomy. During the DWH oil spill, omics’ analyses provided new insights into the degradation capabilities of marine microbes, revealing the dominant pathways that were actively expressed *in situ* in the deep-sea plume (e.g. Mason et al. 2012; Rivers et al. 2013). However, unlike cultivation-based studies, except for single cell genomics analyses presented in Mason et al. (2012 and 2014), omics analyses did not link physiology and identity given the lack of assembled genomes from this data. Using genome assembly, phylogeny, and read mapping, we attempted to bridge the gap and link function and identity of hydrocarbon degraders indigenous to the deep-sea in the GOM and in the global ocean.

Consistent with 16S rRNA based studies, the Gammaprotobacteria that were reported as being abundant members of the plume microbial community, such as *Oceanospirillales*, *Colwellia* and *Cycloclasticus* (Hazen et al. 2010; Valentine et al. 2010; Kessler et al. 2011; Mason et al. 2012; Redmond and Valentine 2012; Kleindienst et al. 2016) were captured in the assembled genomes. We would note that the 16S RNA gene in DWH *Bermanella* MAG BM58_1, the most abundant microbe in our analyses, was 99.93% similar to DWH *Oceanospirillales* clones BM580104 (HM587889.1) and 99.33% similar to DWH *Oceanospirillales* clone OV01102/03 (HM587890.1) (Hazen et al. 2010). This MAG was also 98.36% similar to the dominant *Oceanospirillales* in the plume 16S rRNA gene pyrotag data (Mason et al. 2012). In addition to the dominant players in the plume, we also assembled Gammaproteobacteria classified as UBA11654 and representatives of the SAR324 phylum, neither of which have been previously recognized as members of the plume community.

DWH MAGs were most similar to other DWH MAGs within the same clades, however, there were several notable exceptions. For example, using a different assembly method than we used, the DWH O. Desum v2 genome was highly similar to DWH MAGs (*Bermanella*), as were some MAGs described in Hu et al., (2017), and, in the case of *Colwellia*, isolates such as *Colwellia psychrerythraea 34H*, which was cultured from Artic sea ice (Methé et al., 2005). Additionally, SAR324 DWH MAGs were closely related to SAR324 cluster bacterium SCGCAAA001-C10 from the South Atlantic at 800 m depth (Swan et al., 2011) and to SAR324 cluster bacterium SAR324-CTD7B sampled from a hydrothermal plume in the Atlantic (Cao et al., 2016) (Fig 1). The relatedness, which in some cases was high, of these DWH MAGs to microbes sampled from a broad geographic distribution suggested that the genomic content encoded in the DHW MAGs may be more broadly representative of metabolic schemes employed by microbes in the global ocean, rather than just those residing in the deep-sea in the Gulf of Mexico. These additional genomes are medium to high quality (Table 1) which is particularly important if our MAGs were less than complete, as was the case with DWH *Colwellia*.

Microbial abundances and transcriptional activity as determined by MG and MT read mapping to genomes were consistent with the previous studies that evaluated microbial abundances in the plume (Mason et al. 2012; Redmond and Valentine 2012; Dubinsky et al. 2103; Rivers et al. 2013), in that *Bermanella* was both the most abundant and active microbe in late May 2010. Read recruitment also revealed that while *Cycloclasticus* and *Colwellia* genomes recruited MG reads, RPKM values were much lower than for *Bermanella* genomes and only low levels of MT reads were recruited to these genomes (Fig 2). In agreement with our findings, *Cycloclasticus* and *Colwellia* were reported to be present early in the spill history, but these clades were more abundant, and perhaps more active later in the DWH spill (e.g. Redmond and Valentine 2012). UBA11654 and SAR324 were not reported as being members of the plume community, yet their genomes did in fact recruit MG and MT reads, albeit at lower levels than the aforementioned clades, suggesting a previously unrecognized role for members of these clades in hydrocarbon consumption.

All 38 genomes analyzed encoded hydrocarbon degradation, despite the fact that the majority of the non-DWH plume microbes included in our analyses were sampled from non-hydrocarbon contaminated environments that geographically spanned the global ocean and was inclusive of the surface to deeper in the water column, and marine sediments. The ability to degrade aliphatic and aromatic hydrocarbons was observed, with some pathways that appeared to be clade specific. For example, all *Bermanella* genomes coded for non-gaseous n-alkane degradation, all *Cycloclasticus* genomes coded for methane oxidation, and all SAR324 and all but one UBA11654 genomes coded for aromatic hydrocarbon degradation (Fig 4). In contrast, long chain n-alkanes and/or cyclic alkane degradation was a ubiquitous pathway encoded in members of all clades except SAR324. Read recruitment to these 38 genomes revealed that during the time our samples were obtained in late May-early June 2010, non-gaseous n-alkane degradation was the dominant pathway, while gaseous n-alkane and long chain n-alkane/cycloalkane degradation genes were expressed, but recruited fewer MT reads. Although MG reads coding for aromatic degradation were recruited to the genomes encoding this pathway, particularly in the uncontaminated sample collected from plume depth, no genes were expressed.

During the DWH oil spill, methane was one of the most abundant hydrocarbons input to the GOM. Oxidation rate measurements showed that methane oxidation was an active microbial process (Crespo-Medina et al., 2014; Kessler et al., 2011), while omics studies reported an abundance of pmo transcripts early in the spill history (Mason et al. 2012; Rivers et al. 2013). Taken together the chemical and biological data suggested that methane oxidation was an important process, yet the identity of the microorganisms carrying out this process remained enigmatic. Our analyses revealed that methane oxidation was encoded in all *Cycloclasticus*, and in one *Colwellia*, which provided an important, yet missing link between function and identity.

Methane oxidation is largely undocumented in *Cycloclasticus*. For example, despite its close phylogenetic relationship with canonical methanotrophs (*Methylobacter* and *Methylomonas*) *Cycloclasticus pugetii*, a well-known aromatic hydrocarbon (naphthalene, phenanthrene, anthracene and toluene) degrading bacteria isolated from marine sediments, is not able to use methane for carbon (Dyksterhouse et al. 1995). Using stable isotope probing (Redmond & Valentine, 2012) revealed that indigenous deep-sea *Cycloclasticus* sampled in the GOM deep-sea plume are capable of degrading gases as shown by incorporation of ^13^C labelled propane and ethane, but not methane. Similarly genomic and transcriptomic analyses of *Cycloclasticus* symbionts of mussels and sponges that reside at deep-sea oil and gas seeps can use propane, ethane and butane, but not methane (Rubin-Blum et al., 2017). Although *Cycloclasticus* is not known to degrade methane, GTDB taxonomy which, is based on the alignment of single-copy marker proteins, previously led to the *Cycloclasticus* clade being reclassified as a fourth family in the *Methylococcales* order (Orata et al. 2018). Given that all *Cycloclasticus* encoded *pmoABC* genes, suggesting a methanotrophic lifestyle, our results support the reclassification to the *Methylococcales*, which are a clade of canonical methanotrophs. Additionally, protein tree analyses supported the close phylogenetic relationships between DWH MAGs presented herein and two MAGs (*Cycloclasticus sp. Symbiont of Poecilosclerida sp.* And *Cycloclasticus sp. Symbiont of Bathymodiolus heckerae specimen P*, see Supp. Fig 1) from (Rubin-Blum et al. 2019). Taken together the data suggested that, like other *Methylococcales*, the *Cycloclasticus* analyzed appear to be capable of gaseous hydrocarbon degradation, including methane.

*Colwellia* that were capable of consuming gases were reported in Mason et al. (2014) through SAG assembly, gene annotation and using SIP (Redmond and Valentine 2012). In Mason et al. (2014) the *Colwellia* SAG was hypothesized to have a butane monooxygenase, which is known to preferentially uptake gases other than methane. Using SIP Redmond and Valentine (2012) reported that *Colwellia* was consuming ethane and propane, but not methane. Here, we found evidence that extends *Colwellia*’s gaseous hydrocarbon degradation capabilities to include methane oxidation.

Taken together, genome assembly and read mapping definitively links methane oxidation with taxonomy. Specifically, *Cycloclasticus* and a *Colwellia* were capable of consuming methane, an active process carried out early in the DWH spill history. In fact, based on MG read recruitment from the uncontaminated sample collected from plume depth, methane oxidation by indigenous *Cycloclasticus* and to a lesser degree *Colwellia* appears to be an abundant hydrocarbon degradation metabolism in indigenous microbes residing in the deep-sea in the GOM, albeit expression levels were lower than other hydrocarbon degradation pathways. This suggests that methane oxidation is mediated by previously unrecognized microbes, that in the case of *Colwellia* and until recently, *Cycloclasticus*, are outside of the canonical methanotrophic clades that are typically thought to carry out this reaction. This is similar to the recent findings of active, non-canonical methanotrophs that were reported in the GOM (Howe et al. 2023). Further, the fact that all *Cycloclasticus* encoded methane oxidation, regardless of where it was sampled from, further supports its taxonomic placement as a family in the *Methylococcales*, a microbial group that has become a paradigm of marine methane oxidation.

Non-gaseous n-alkane degradation was the most abundant hydrocarbon degradation pathway in terms of MG and MT read recruitment at the time we sampled, which is consistent with previous reports that suggested non-gaseous n-alkane degradation was the most abundant metabolic strategy employed be indigenous microbes to degrade DWH oil in the deep sea plume (Mason et al. 2012; Rivers et al. 2013). Degradation of medium chain n-alkanes (C_5_-C_26_) was largely attributed to *Bermanella* that encoded *alkB* (Mason et al. 2012). While *alkB* was encoded in the genomes of all non-DWH *Bermanella*, two DWH *Colwellia* MAGs, one non-DWH *Cycloclasticus*, and one UBA11654 (Fig 4), none of the DWH *Bermanella* encoded this gene. Instead, the DWH *Bermanella* in the plume encoded and expressed CYP153. We also identified rubredoxin as present and expressed in all DWH *Bermanella* MAGs, *Bermanella marisrubri RED65*, and *Bermanella sp. CPC64*. In *A. borkumensis* rubredoxin is part of the complete AlkB operon, and is the second step in alkane oxidation (Sabirova et al. 2006). Rubredoxins are integral electron transfer components necessary for alkane hydroxylation by AlkB (Nie et al. 2014; Van Beilen et al. 2002). It has been suggested that it could theoretically serve the same function with CYP153 (Nie et al. 2014). The presence of CYP153 monooxygenase and rubredoxin genes in these genomes suggested that n-alkane degradation proceeds from CYP153 to rubredoxin, which would represent the two domain architecture for n-alkane degradation (Nie et al. 2014), is a likely pathway used by the dominant microbes to degrade oil in the deep-sea consortium.

Additionally, three DWH *Bermanella* MAGs that encoded CYP153 also encoded cytochrome P450, ferredoxin, and ferredoxin reductase suggesting a three domain CYP153 architecture for n-alkane degradation (Nie et al. 2014). Further, if rubredoxin functions similarly when paired with CYP153 as it does with AlkB this would indicate that DWH *Bermanella* used two pathways to degrade non-gaseous n-alkanes. The fact that DWH *Bermanella* encoded CYP153 rather than AlkB as originally proposed (e.g. Mason et al. 2012) may be due to the high abundance of C_8-_C_16_ n-alkanes compared to longer chain n-alkanes in Macondo oil that has been reported (e.g. see Figure 5 in Kujawinski et al. 2020). This hypothesis is supported by heterologous expression experiments with *A. dieselolei* AlkB, CYP153, and AlmA (discussed below) in Liu et al. (2011). They showed that CYP153 was upregulated in the presence of C_8_-C_16_ n-alkanes, while AlkB responded to C_12_-C_26_. The more focused range of n-alkanes that led to upregulation of CYP153 compared to AlkB reported by Liu et al. (2011) may suggest that CYP153 is an enzyme that is more fine-tuned to C_8-_C_16_ n-alkane degradation, which is consistent with the profile of Macondo oil. Thus, degradation of non-gaseous n-alkane hydrocarbons using multiple, previously undescribed pathways by DWH *Bermanella*, likely gave this microbe an advantage in that the dominant n-alkanes it degrades were the most abundant in Macondo oil. This enzyme’s fine-tuning is a plausible explanation for dominance of *Bermanella* in the deep-sea plume.

Long-chain (>C_18_) n-alkane/cyclic degradation was a ubiquitous pathway encoded in all clades analyzed here, except SAR324. The 4-hydroxyacetophenone monooxygenase (HAPMO)/flavin-containing monooxygenase identified in DWH *Bermanella* MAGs herein were highly abundant and expressed in the plume samples and less abundant in the uncontaminated sample (Fig 4). HAPMO is a flavin-containing monooxygenase that catalyzes NADPH and oxygen-dependent Baeyer-Villiger oxidation of 4-hydroxyacetophenone. It is related to several BVMOs including cyclohexanone monooxygenase and *almA*. These HAPMO genes had ∼30% similarity with *almA* genes and ∼24% similarity with *A. borkumensis’* cyclohexanone monooxygenase. Other flavin-containing monooxygenases, including *almA*, were annotated in DWH and non-DWH *Cycloclasticus* and UBA11654. These genes were most abundant in the uncontaminated sample, compared to the low abundances in plume samples (Fig 4). None of these genes were expressed (Fig 4).

(Maeng et al. 1996) identified a flavin-containing n-alkane dioxygenase in *Acinetobacter sp.* Strain M-1 which allowed for growth on medium to long-chain n-alkanes C_10_ to C_30_. Later, (Throne-Holst et al. 2007) reported that AlmA in *Acinetobacter sp. DSM 17874* supports its growth on long-chain alkanes up to C_36_. Liu et al. (2011) carried out heterologous expression experiments and reported that AlmA in in the marine hydrocarbon degrader, *A. dieselolei,* functions in long-chain n-alkane degradation. They further reported that AlmA was upregulated in the presence of long chain n-alkanes ranging from C_22_ to C_36_. Putative *almA* genes have also been identified in *A. borkumensis* SK2 (Rojo 2009) which degrades long change n-alkanes up to C_32_. Further, AlmA was recently identified in several *Oceanospirillales* MAGs that were assembled from the Mariana Trench (J. Liu et al. 2019).

Although less abundant than medium chain n-alkanes, long chain n-alkanes up to C_38_ are constituents of Macondo oil (Kujawinski et al. 2020). Thus, the high abundance of flavin-containing monooxygenases in DWH *Bermanella, Cycloclasticus* and UBA11654 MAGs, with varying degrees of similarity with *almA* (30 to > 60% similarity), that were expressed in the plume, suggested that oxidation of long-chain n-alkanes up to C_36_ and perhaps longer chain alkanes, was in important microbial degradation strategy used by indigenous microbes during the DWH oil spill. One that has not yet been recognized. Further, non-DWH *Bermanella, Cycloclasticus* and UBA11654 MAGs from microbes sampled globally, also encoded flavin-containing monooxygenases, including *almA*, suggesting long-chain n-alkane degradation is a ubiquitous pathway in hydrocarbon degrading microbes in the global ocean.

Cyclohexanone monooxygenase, a BVMO, was identified through annotation in DWH and non-DWH *Cycloclasticus* and non-DWH UBA11654. This gene annotation suggested cyclic alkane degradation, yet an enzyme, such as cyclohexane monooxygenase that initiates cyclic alkane degradation, was not identified in any MAG. However, AlkB and cyclohexanone monooxygenase were identified in UBA11654

*Methylococcaceae bacterium TMED69*, sampled from the photic zone in the Mediterranean Sea (Tully et al. 2018a) and in *Cycloclasticus Methylococcaceae bacterium UBA975* from North Sea water (Parks et al. 2017). It was postulated that cyclic alkane degradation by *A. borkumensis* is initiated with AlkB, with the second step being mediated by cyclohexanone monooxygenase, allowing for degradation of both linear and cyclic alkanes (Sabirova et al. 2006). DWH *Cycloclasticus* cyclohexanone monooxygenase genes were more abundant in the uncontaminated sample compared to the plume, but were not expressed. Mason et al. (2012) obtained an *Oceanospirillales* (*Bermanella*) single-cell genome that encoded cyclohexanone monooxygenase, while none of the *Bermanella* analyzed herein encoded that gene. The DWH *Bermanella* HAPMO/flavin-containing genes may be involved in cyclic alkane degradation, but the data suggested that an enzyme other than cyclohexanone monooxygenase is involved.

Members of all clades encoded for aromatic hydrocarbon degradation genes, although only one non-DWH *Bermanella* was represented. Of these clades, all members of SAR324 coded for aromatic hydrocarbon degradation, but no other hydrocarbon degradation pathway was observed in this group. Only representatives of DWH and non-DWH members of *Colwellia*, UBA11654, and SAR324 recruited MG reads to these genes. BTEX was abundant in the plume (Reddy et al. 2011), yet none of the genes involved in aromatic hydrocarbon degradation identified here recruited any transcripts. This lack of transcript recruitment suggested that aromatic hydrocarbon degradation was not an active process carried out by the microbes presented herein in the GOM water column one month after the spill began (Fig 4). Aromatics such as naphthalene (a PAH and known carcinogen) surfaced and was thought to evaporate rapidly (Ryerson et al. 2011). Although active aromatic hydrocarbon degrading microbes were not observed in the DWH plume, the ability to degrade these hydrocarbons appears to be global, with this pathway identified in samples from other parts of the GOM, the North Atlantic, the Red Sea, the North Sea, and from sediments and crustal fluids.

Beyond hydrocarbon substrate availability, the bioavailability of nutrients, including nitrogen and phosphorus, is a key factor in the speed and success of hydrocarbon degradation, therefore, the abundance and expression of nutrient acquisition and transport genes can also provide insight into the microbial hydrocarbon degradation process (Head et al. 2006). DWH *Bermanella* sulfur transport, phosphate/phosphonate transport, and APase genes were highly abundant and expressed, but other genes involved in sulfate reduction and phosphorus cycling were not abundant or expressed by members of any clade (Fig 5).

Most *Colwellia* encoded for complete denitrification, a pathway which has previously been shown in members of this clade (Methé et al. 2005; Mason et al. 2014), while members of other clades encoded for some steps but not the entire pathway (Fig 5). Presence and expression of genes involved in anaerobic respiration indicate alternate electron acceptors necessary to continue hydrocarbon degradation in the oxygen anomaly which was created due to the high respiration of hydrocarbons and nutrients by the bloom of microorganisms (Dubinsky et al. 2013; Rivers et al. 2013). Although there was no overall significant oxygen depletion inside or outside the plume, nitrate concentrations were significantly lower in the plume (Hazen et al. 2010). Therefore, denitrification by *Colwellia* is one possible explanation for the difference in nitrate concentrations inside and outside of the plume.

The abundance and expression of siderophore biosynthesis genes from DWH and non-DWH *Bermanella* and *Colwellia,* and to a lesser extent in *Cycloclasticus*, in these samples indicates the ability to continue, and potentially increase, hydrocarbon degradation in an iron-depleted environment (Lu et al. 2012). Further, some *Colwellia* lacked genes coding for siderophore biosynthesis, but had the ability to transport siderophores, which is consistent with *Colwellia* scavenging siderophores (Methé et al. 2005; Mason et al. 2014). Iron-depletion in the plume was reported (Joung & Shiller, 2013), and likely arose from the iron demand of *Bermanella*, *Colwellia,* and *Cycloclasticus* that were actively carrying out hydrocarbon degradation. However, it is unlikely that iron depletion reached concentrations that presented a growth-limiting factor (Joung & Shiller, 2013).

All *Bermanella, Colwellia*, and *Cycloclasticus*, but only one SAR324 and no UBA11654, encoded methyl-accepting chemotaxis proteins (MCPs), with highest expression by members of *Bermanella* (Fig 5). Some MCPs have been found to aid in sensing alkanes and other substrates and therefore may be linked with hydrocarbon degradation (Smits et al. 2003; Mason et al. 2012; Wang and Shao 2013, 2014; Liu et al. 2019). While all clades encoded flagellar biosynthesis and motor genes involved in motility, only 3/9 UBA11654 coded for motility (Fig 5). *Cycloclasticus* motility genes were some of the most abundant and expressed across all samples. This clade has long been known to contain motile members (Dyksterhouse et al. 1995; Wang et al. 1996) and may have been able to move towards degradable hydrocarbon substrates.

All clades encoded cold-shock proteins (Csps), such as CspA, which was one of the most highly expressed genes in our study. Csps are expressed in response to rapid temperature decrease (cold shock), as well as a wider range of stressors including those related to pH, starvation, and oxidative tolerance (Keto-Timonen et al. 2016). During stress but specifically cold shock, cell membrane fluidity and enzyme activity decrease, leading to reduced efficiency of transcription and translation due to formation of nucleic acid secondary structures (Keto-Timonen et al. 2016). Csps act as nucleic acid chaperons that may prevent these secondary structures from forming at these low temperatures or other less than optimal conditions (Keto-Timonen et al. 2016). Thus, high levels of CspA gene expression indicated that plume microbes were attempting to counteract the decrease in efficiency in transcription and translation, facilitating continued hydrocarbon degradation in the low temperature deep-sea plume. The *Colwellia* clade contains known psychrophiles (Methé et al. 2005; Redmond & Valentine, 2012; Hamdan, 2018) and while the temperature of these deep-sea plume samples was around 4°C, these cold-shock genes may also play a wider role in stress tolerance (Keto-Timonen et al. 2016 and references therein). Cold-shock genes encoded and expressed in these samples supports their advantageous involvement in response to hydrocarbon inputs (Mason et al. 2014; Liu et al. 2019), allowing microbial consortia to sense, respond and bloom, while cycling nutrients and hydrocarbons at depth.

The DWH pangenome, or core genes, included genes and pathways that are involved in hydrocarbon degradation. For example, citrate synthase is involved in the TCA/glyoxylate pathway, which can be a final step of aromatic hydrocarbon degradation or can take place after fatty acid beta oxidation (Rivers et al. 2013). Aldehyde dehydrogenase can facilitate steps in gaseous (Supp. Fig 2) and non-gaseous n-alkane oxidation, and conversion of fatty aldehydes to fatty acids for incorporation into the beta oxidation pathway (Rivers et al., 2013b; Rojo, 2009a). Oxidation of alkanes results in formation of fatty acids which are degraded in the beta oxidation pathway (Rojo 2009). Further, the pangenome included multiple steps of the beta oxidation pathway which is the primary metabolism pathway for fatty acids derived from degradation of alkanes and aromatics (Rivers et al. 2013). Rivers et al. (2013) showed that beta-oxidation was one of the main pathways in which alkanes and aromatics were metabolized. Here, these transcripts were either enriched or significantly enriched as were transcripts of genes involved in the initial hydrocarbon oxidation steps. These core pangenome genes suggested that marine microbial hydrocarbon degradation is broadly distributed across multiple microbial clades. Linking cross-clade hydrocarbon degradation core genes with clade-specific hydrocarbon degradation genes/pathways discussed above, provided missing information that identifies both the microbe and pathway used in degrading specific hydrocarbon substrates. These findings help resolve previous studies that linked hydrocarbon degradation to the dominance and succession of specific microbial groups over the course of the DWH oil spill (Hazen et al., 2010; Valentine et al., 2010; Kessler et al., 2011; Redmond & Valentine, 2012; Dubinsky et al., 2013).

## Conclusion

Collectively the microbes analyzed in our study spanned the global ocean and included the surface ocean to the deep-sea in the water column, sea ice, crustal fluids and marine sediments. Unifying all 38 microbes was the ability to degrade hydrocarbons, which was unexpected given the fact that many were sampled in non-hydrocarbon contaminated marine environments. This suggested that hydrocarbon degradation capabilities are maintained in the microbial population, even in the seeming absence of oil and gas substrates. Given that marine microbial hydrocarbon degradation is an essential ecosystem service, beyond cultivation studies, the missing linkages between taxonomy and function has left a critical knowledge gap regarding the intrinsic bioremediation capabilities of uncultured, indigenous microbes. Here we provide these missing linkages and reveal novel hydrocarbon degradation strategies not previously attributed to specific clades. Further, our analyses revealed that the microbes that dominated the deep-sea plume possessed enzymes and pathways that were fine-tuned to Macondo oil. We also report that beyond the chemical composition of Macondo oil, the ability to sense and move to degradable hydrocarbon substrates likely played a significant role in determining what microbes dominated the deep-sea plume. Finally, in addition to natural and anthropogenic inputs of hydrocarbons into the marine environment due to oil extraction and transportation activities, petroleum-based pollution in the form of plastics is also a major issue. Plastics enter the ocean at approximately 10 million metric tons annually (Jambeck et al., 2015), providing surfaces for microbial colonization, while also leaching hydrocarbons into the surrounding environment (reviewed in Muriel-Millán et al., 2021; Pinto et al., 2022). Thus, plastic pollution contributes to the pool of hydrocarbons available for microbial consumption. Microbes capable of alkane degradation, including *Bermanella*, have been shown to colonize plastic particles and/or consume plastic leachates (Muriel-Millán et al., 2021; Stubbins et al., 2021; Wright et al., 2021; Cao et al., 2022). Collectively, the results of our analysis of multi-omics data, revealed the identity and metabolic strategies of uncultured hydrocarbon degraders in the Gulf of Mexico and in the global ocean. This new information allows for better prediction of the microbial response to hydrocarbon pollution, in all its forms, in the marine environment, and ultimately the ability to predict the ecological consequences of such inputs.

## Acknowledgements

We thank the captain, crew and science teams aboard the R/V Ocean Veritas and the R/V Brooks McCall. This work was supported by a subcontract from the University of California at Berkeley, Energy Biosciences Institute to Lawrence Berkeley National Laboratory under its US Department of Energy contract DE-AC02-05CH11231 and by a grant from The Gulf of Mexico Research Initiative RFP-VI: Research Grants [OUM Consortium for Simulation of Oil-Microbial Interactions in the Ocean (CSOMIO)].

## Data Availability Statement

MG reads are publicly available in NCBI (PRJNA336903-05) in IMG Gold Study ID Gs0063184 and on the Mason server http://mason.eoas.fsu.edu in the DWH_plume directory. Amplified cDNA reads are publicly available in NCBI (PRJNA839076). MAGs are also available in NCBI under SAMN37094393-95, SAMN37094224-28, SAMN37094233-36 (accession numbers have been requested for each MAG) and are linked with their respective metagenome sequence data in NCBI (PRJNA336903-05). These MAGs will be made publicly available in NCBI upon manuscript acceptance.

